# Lung injury induces a polarized immune response by self antigen-specific Foxp3^+^ regulatory T cells

**DOI:** 10.1101/2023.02.09.527896

**Authors:** Daniel S. Shin, Sneha Ratnapriya, Creel Ng Cashin, Lucy F. Kuhn, Rod A. Rahimi, Robert M. Anthony, James J. Moon

**Affiliations:** Center for Immunology and Inflammatory Diseases, Division of Rheumatology, Allergy, and Immunology, Massachusetts General Hospital, Charlestown, MA, USA; Division of Immunology, Boston Children’s Hospital, Boston, MA, USA; Division of Pulmonary and Critical Care Medicine, Massachusetts General Hospital, Boston, MA, USA; Harvard Medical School, Boston, MA, USA

**Keywords:** CD4^+^ T cell, peripheral tolerance, self antigen, Foxp3, regulatory T cell, Tregs, T cell repertoire, autoimmunity, epitope spreading, peptide:MHC tetramer

## Abstract

Self antigen-specific T cells are prevalent in the mature adaptive immune system, but are regulated through multiple mechanisms of tolerance. However, inflammatory conditions such as tissue injury may provide these T cells with an opportunity to break tolerance and trigger autoimmunity. To understand how the T cell repertoire responds to the presentation of self antigen under highly stimulatory conditions, we used peptide:MHCII tetramers to track the behavior of endogenous CD4^+^ T cells with specificity to a lung-expressed self antigen in mouse models of immune-mediated lung injury. Acute injury resulted in the exclusive expansion of regulatory T cells (Tregs) that was dependent on self antigen recognition and IL-2. Conversely, conventional T cells of the same self antigen specificity remained unresponsive, even following Treg ablation. Thus, the self antigen-specific T cell repertoire is poised to serve a regulatory function during acute tissue damage to limit further damage and the possibility of autoimmunity.

## Introduction

The adaptive immune system contains a vast repertoire of T cells expressing unique T cell receptor (TCR) specificities that can respond to virtually any foreign antigen. During development, thymocytes expressing TCRs that recognize self antigens are normally deleted or programmed into regulatory T cells (Tregs) under mechanisms of central tolerance. However, this process is not absolute, and many conventional self antigen-specific T cells (Tconv) mature and populate the peripheral immune repertoire^1–5^, where multiple overlapping mechanisms of peripheral tolerance restrain them from potentially pathogenic fates^6^. In experimental systems, compromising the integrity of steady-state peripheral tolerance can result in the activation of these cells and lead to autoimmunity^7^. More recently, the use of immune checkpoint inhibitors in cancer therapies has provided direct evidence in humans that the impairment of peripheral tolerance, whether by blocking inhibitory signals in activated T cells or impeding Treg activity, can cause self antigen-specific T cells to attack healthy tissue and drive the immune-related adverse events associated with these treatments^8–11^.

While many factors contribute to the pathogenesis of autoimmune diseases, there is often circumstantial evidence linking inflammatory events to the initiation of disease, and many experimental models are indeed designed around the deliberate priming of immune responses against a self antigen^12–16^. Thus, characterizing how steady-state peripheral tolerance fails under conditions of inflammation may provide crucial insights into how the activation of self antigen-specific T cells occurs during the early stages of autoimmunity. One such process, known as “antigen or epitope spreading”, describes a cascade of events wherein the inflammation from an initial immune response causes nearby self antigens to be secondarily processed by activated antigen presenting cells (APCs) and presented to self antigen-specific T cells, leading to subsequent autoreactive immune responses^17^. Epitope spreading has been observed in several autoimmune diseases^18–21^ and is well characterized in mouse models of relapsing-remitting experimental autoimmune encephalomyelitis (EAE) ^22–24^. This mechanism of activation contrasts with the phenomenon of bystander activation in which self-reactive T cells are activated by an inflammatory environment in an antigen-independent manner^25^.

Unfortunately, studying the physiologic response of self antigen-specific T cells upon recognition of their cognate antigen during inflammation is technically challenging due to the extremely low frequencies of T cells specific for any given antigen^26^, let alone a self antigen for which many or most T cells may have been clonally deleted. Studies relying on the use of TCR-transgenic T cells either adoptively transferred into, or crossed onto, host mice expressing the cognate antigen have informed much of our knowledge regarding peripheral tolerance^27^, but due to the highly manipulated nature of these experimental systems, they cannot be expected to fully recapitulate how a true self antigen-specific T cell population under steady-state regulation responds to the acute presentation of cognate antigen under inflammatory conditions, and what mechanisms of tolerance are most relevant in these cases.

Our lab has developed the use of peptide:MHC tetramer reagents and model antigen-bearing mice to rigorously analyze how endogenous self antigen-specific T cell populations are regulated in the steady state in a tissue specific manner^2,28^. Here, we leveraged the advantages of our experimental systems to directly address how immune-mediated lung injury in these mice leads to activation of endogenous self antigenspecific CD4^+^ T cells. We found that, contrary to direct priming, activation by epitope spreading leads to near exclusive expansion of self antigen-specific Tregs over their Tconv counterparts, suggesting that the physiologic response of the self antigenspecific CD4^+^ T cell repertoire during acute tissue injury is to expand Tregs which regulate ongoing autoimmune activity^29^.

## Results

### Autoimmune T cell-mediated tissue injury leads to expansion of self antigenspecific Tregs

We have previously reported the generation of a conditional transgenic mouse which, in the presence of Cre recombinase, expresses a Universal Self Antigen (USA) consisting of two model I-A^b^-restricted CD4^+^ T cell epitopes, 2W and gp66, embedded within a membrane-bound form of chicken ovalbumin (OVA) ^28^. Crossing these *USA* mice to *CC10-Cre* transgenic mice results in USA antigen expression restricted to lung epithelial cells via the Clara cell (CC10) promoter. We can directly track and characterize lung self antigen-specific CD4^+^ T cells in the mature peripheral repertoire of these *CC10-Cre* x *USA* mice (*CC10-USA*) via the use of 2W:I-A^b^, gp66:I-A^b^, or OVA:I-A^b^ tetramers^28,30^. As reported earlier, naïve 2W:I-A^b^-specific T cells are present in reduced numbers compared to wildtype (*WT*) littermates, slightly enriched for Foxp3 expression, and absent in the lung parenchyma, demonstrating the effects of steady state tolerance on this self antigen-specific T cell population (data not shown).

To investigate how the self antigen-specific T cell repertoire responds to presentation of cognate antigen during acute autoimmune tissue injury, we adoptively transferred congenically marked gp66:I-A^b^-specific CD4^+^ T cells isolated from *Smarta* TCR transgenic mice into *CC10-USA* mice and then analyzed endogenous 2W:I-A^b^-specific CD4^+^ T cells in the secondary lymphoid organs (SLO) and lungs 7 days later (Fig. 1A, S1A). As expected, the transferred *Smarta* CD4^+^ T cells accumulated at higher frequencies in the SLO and lungs of *CC10-USA* mice compared to *WT* littermates (Fig. S1B-C). Analysis of bronchoalveolar lavage (BAL) fluid revealed an increase in IgM levels that is consistent with leakage from epithelial tissue damage (Fig. S1D). We did note a small subset of the *Smarta* CD4^+^ T cells transferred into *CC10-USA* mice expressed Foxp3, indicating a level of auto-regulation of these deleterious T cells (Fig. S1C).

**Figure 1.**
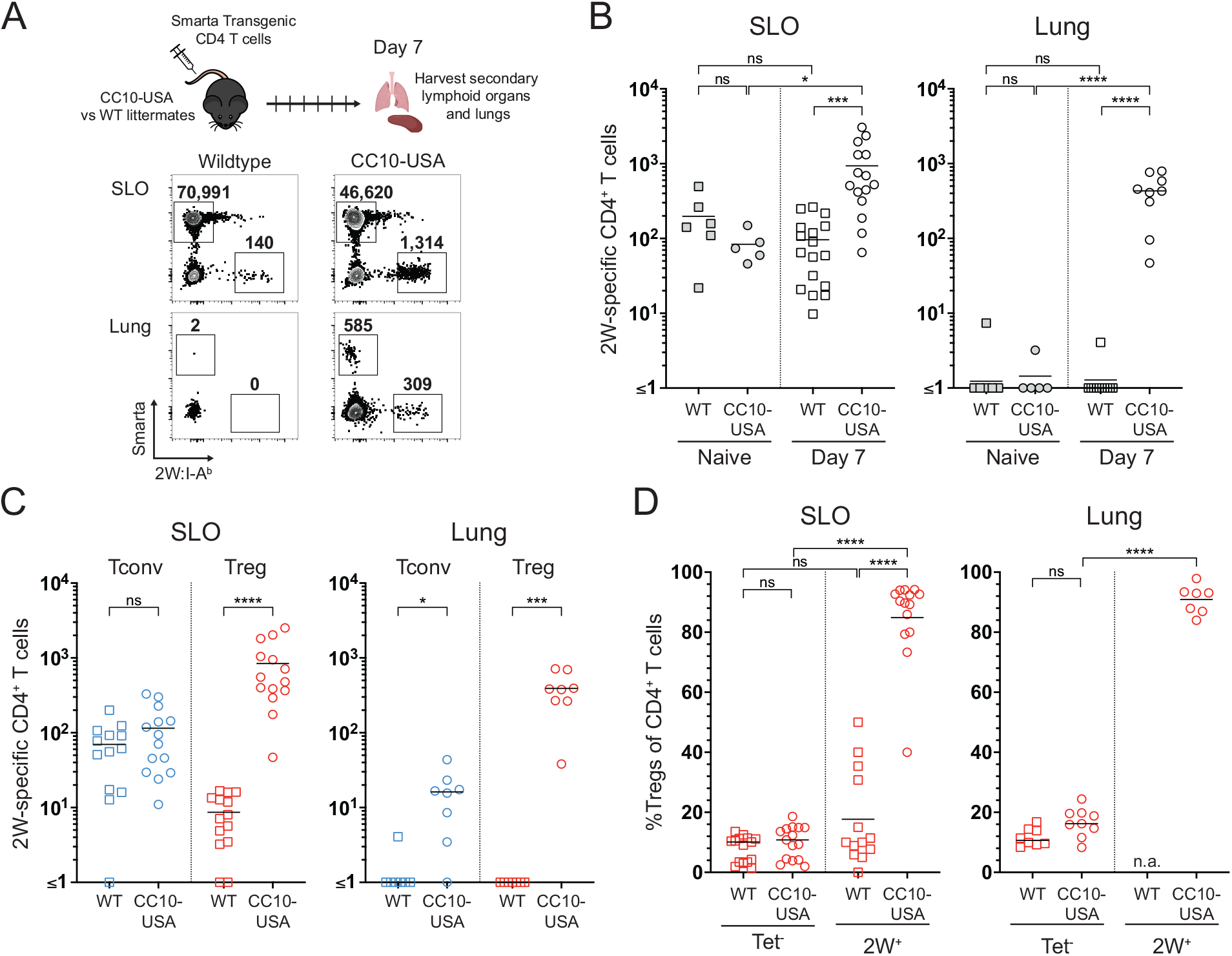
Asymmetric expansion of Foxp3^+^ Tregs within a lung self antigenspecific T cell population following tissue injury. (**A**) gp66:I-A^b^-specific CD4^+^ T cells from *Smarta* mice were adoptively transferred into *CC10-USA* mice or wildtype (*WT*) littermate controls, and 7 days later, the mice were euthanized for tissue harvest. Representative flow cytometry plots of CD4^+^ T cells following magnetic bead-based enrichment of pooled spleen and lymph nodes (SLO) or lungs. Numbers above gates indicate total number of 2W:I-A^b^-specific or *Smarta* CD4^+^ T cells calculated for the whole mouse. (**B**) Quantification of total 2W:I-A^b^-specific CD4^+^ T cell numbers in the SLO and lungs of *CC10-USA* mice 7 days following *Smarta* transfer. (**C**) Quantitative breakdown of 2W:I-A^b^-specific T cell numbers into Foxp3-negative conventional (Tconv) and Foxp3-positive regulatory (Treg) T cell subsets in the SLO and lungs. (**D**) Proportions of Foxp3^+^ Tregs within bulk or 2W:I-A^b^ specific CD4^+^ T cell populations in the SLO and lungs of *CC10-USA* mice vs. *WT* littermates 7 days following *Smarta* transfer. Each datapoint represents an individual mouse. Bars indicate mean values of each population. Statistical significance was calculated via (**B**, **D**) one-way ANOVA with Tukey’s multiple comparisons test or (**C**) unpaired t test; na, not applicable, ns, not significant, *p < 0.05, **p < 0.01, ***p < 0.001, ****p < 0.0001.

Following *Smarta* transfer, we detected an expansion of host 2W:I-A^b^-specific CD4^+^ T cells in *CC10-USA* versus *WT* recipients or mice that did not receive adoptive transfers (Fig. 1B). Notably, the expansion of 2W:I-A^b^-specific T cells occurred almost exclusively within the Foxp3-expressing compartment, resulting in a strong skewing of the self antigen-specific population towards a Treg phenotype (Fig. 1C-D). In contrast, very few type 1 regulatory T cells (Tr1) were seen, as evidenced by IL-10 expressing Foxp3-negative T cells in *CC10-USA* mice crossed to *10BiT*(IL-10) reporter mice (Fig. S1E-F). A similar expansion of 2W:I-A^b^-specific Tregs was seen when OVA-specific CD8^+^ T cells from *OT-I* TCR transgenic mice were transferred into these mice (Fig. S1G). These results indicate that autoimmune damage of lung epithelium results in a heavily polarized Foxp3^+^ Treg response from the self antigen-specific CD4^+^ T cell repertoire.

### The expansion of self antigen-specific Tregs is dependent on TCR and IL-2 signaling

Given the asymmetric expansion of 2W:I-A^b^-specific Treg over Tconv cells, we set out to characterize their activation requirements. To determine if expansion was dependent on TCR recognition of cognate antigen, *Smarta* CD4^+^ T cells were transferred into *CC10-USA* mice in the presence of a monoclonal antibody that binds to 2W:I-A^b^ complexes (clone W6) and blocks their presentation to CD4^+^ T cells^31^. Compared to an isotype control, the blocking antibody significantly decreased 2W:I-A^b^-specific T cell expansion in the SLO and lungs of *Smarta* recipient mice (Fig. 2A). Of note, no overall effects of the antibody were observed on the Tconv cells.

**Figure 2.**
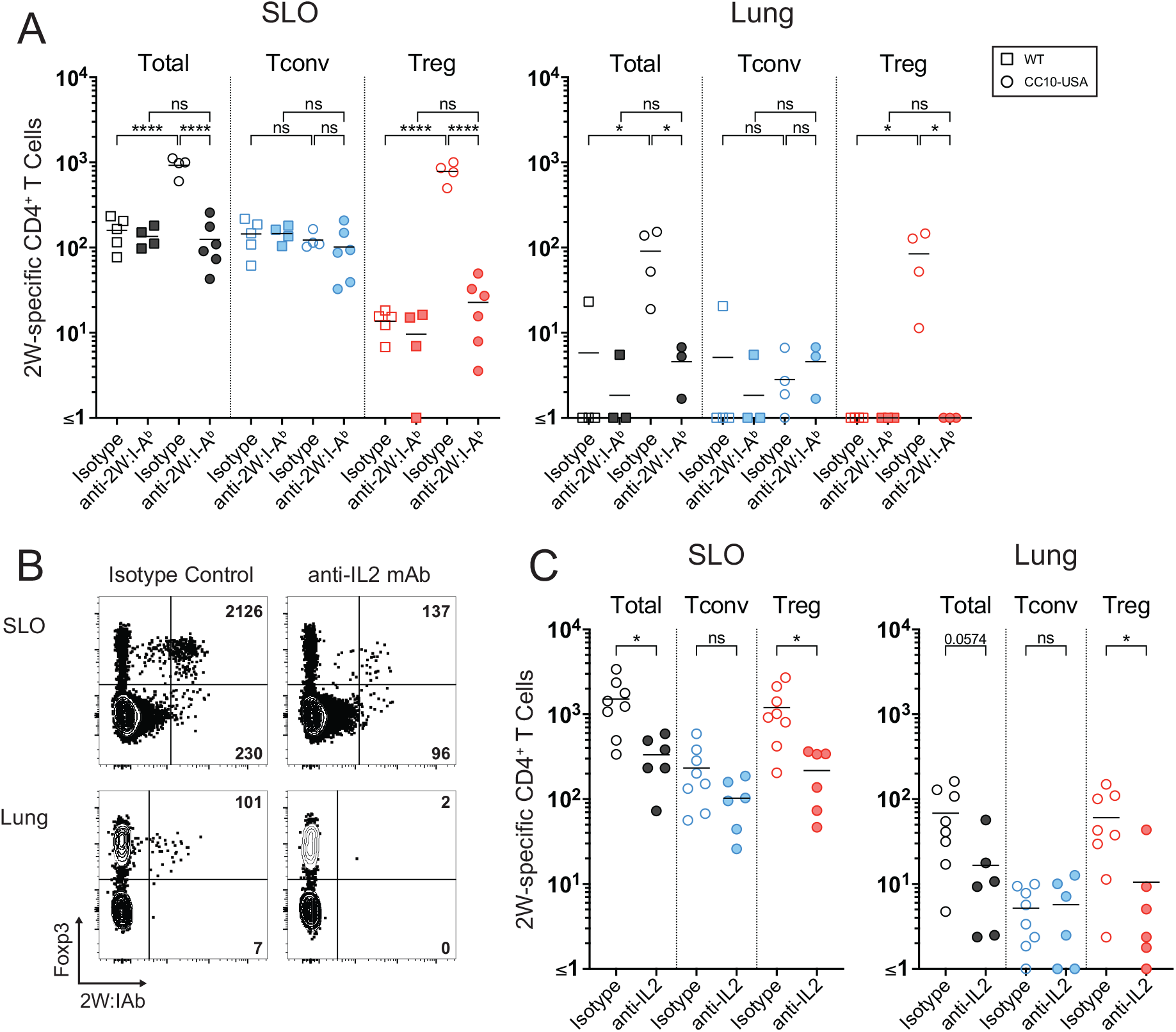
Activation of 2W:I-A^b^ specific Tregs are dependent on the recognition of cognate self antigen and IL-2 signaling. (**A**) 500 μg of 2W:I-A^b^ blocking antibody or isotype control was administered intraperitoneally (*i.p.*) at the time of *Smarta* CD4^+^ T cell transfer into *CC10-USA* mice (circles) and *WT* littermates (squares), and 7 days later, mice were harvested for analysis of 2W:I-A^b^ specific CD4^+^ T cell numbers in the SLO and lungs. Quantitative summary of total 2W:I-A^b^ specific CD4^+^ T cells, further subdivided into Tconvs and Tregs, in the SLO and lungs of each mouse. (**B**) Representative flow cytometry plots of peripheral CD4^+^ T cells in the SLO and lungs 7 days following *i.p*. administration of 500 μg of IL-2 blocking antibody or isotype control at the time of *Smarta* CD4^+^ T cell transfer into *CC10-USA* mice. Numbers within quadrants represent calculated total of 2W:I-A^b^ specific Tconv and Treg populations per mouse. (**C**) Quantification of total 2W:I-A^b^ specific CD4^+^ T cells, further subdivided into Tconvs and Tregs, in the SLO and lungs of *CC10-USA* mice following IL-2 blockade. Each datapoint represents an individual mouse. Bars indicate mean values of each population. Statistical significance was calculated via (**A**) one-way ANOVA with Tukey’s multiple comparisons test or (**C**) unpaired t test; ns, not significant, *p < 0.05, ****p < 0.0001.

We next characterized the cytokine signaling requirements for the expansion of the 2W:I-A^b^-specific Tregs. Interleukin-2 (IL-2) is an important regulator of Treg biology and numerous studies have characterized its role in Treg development, homeostasis, and activation^32^. Additionally, transforming growth factor β (TGFβ) has been shown to be particularly important for the peripheral induction of Tregs, specifically by regulating Foxp3 expression^33^. Lastly, interleukin-10 (IL-10) signaling has been demonstrated to promote Treg identity and function, particularly in mucosal tolerance^34–36^. Following administration of IL-2 blocking antibody into mice during *Smarta* CD4^+^ or *OT-I* CD8^+^ T cells, we found a substantial reduction in the expansion of 2W:I-A^b^-specific T cells in *CC10-USA* mice which was completely attributable to a loss of Treg expansion (Fig. 2B-C). In contrast, we found that blockade of TGFβ or IL-10 had no effect on the expansion of 2W:I-A^b^-specific Tregs following *Smarta* T cell transfer (Fig. S2A-B). Lastly, inhibition of either epitope recognition by TCR or IL-2 signaling, but not IL-10 blockade, led to decreased Tr1 development among 2W:I-A^b^ specific T cells (Fig. S2C-E).

Hence, our results suggest that the 2W:I-A^b^-specific Tregs detected in mice posttransfer are likely to be mature thymically-derived Tregs that expand in an IL-2-dependent manner to the increased presentation of USA self antigen in an inflammatory context.

### Self antigen-specific Tregs are suppressive but not necessary for the regulation of self antigen-specific Tconv cells

To address whether the expanded 2W:I-A^b^-specific Tregs were suppressive, we challenged *CC10-USA* mice 7 days *post-Smarta* T cell transfer with a subcutaneous immunization of 2W peptide emulsified in Complete Freund’s Adjuvant (CFA) and analyzed 2W:I-A^b^-specific T cells from the SLO another 7 days later (Fig. 3A). As shown in Fig. 3B, the expansion of 2W:I-A^b^-specific Tconv cells was reduced in these mice compared to *CC10-USA* mice that had not received *Smarta* T cells. *WT* recipient mouse controls demonstrated that the presence of transferred *Smarta* T cells themselves did not inhibit the expansion of 2W:I-A^b^-specific T cells following immunization. Although *Smarta* T cell transfer into *CC10-USA* mice did not cause a significant increase in the number of 2W:I-A^b^-specific Tregs following immunization, it did result in a substantial decrease in the ratio of Tconv to Tregs in the antigen-specific population, indicating a shift to a less inflammatory immune context (Fig. 3C, right). Thus, the 2W:I-A^b^-specific Tregs that expand following *Smarta*-mediated lung injury are capable of suppressing the activation of their Tconv counterparts.

**Figure 3.**
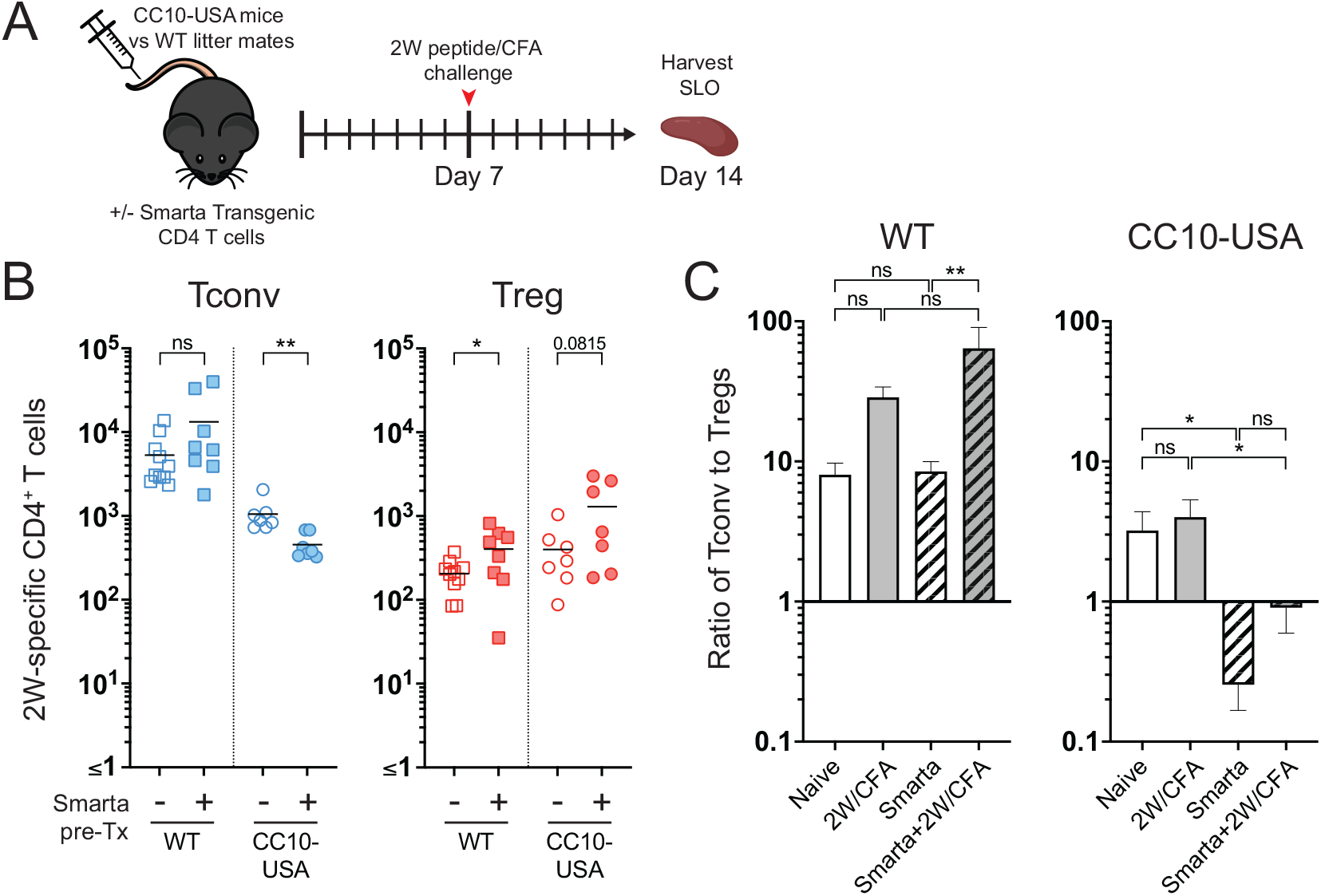
2W:I-A^b^-specific Tregs are able to suppress the expansion of their Tconv counterparts upon peptide immunization. (**A**) Experimental plan to test *in vivo* suppression of 2W:I-A^b^-specific Tconv by expanded 2W:I-A^b^-specific Tregs. Wildtype or *CC10-USA* mice were challenged with 2W peptide + CFA subcutaneously (*s.c*.) 7 days following *Smarta* transfer. Naïve wildtype or *CC10-USA* mice were also immunized as controls. SLO samples were harvested 7 days postimmunization for enumeration of 2W:I-A^b^ specific CD4^+^ T cells. (**B**) Quantification of 2W:I-A^b^ specific CD4^+^ Tconv and Treg numbers in the SLO of *CC10-USA* or *WT* littermate mice with or without prior *Smarta* transfer 7 days after 2W/CFA challenge. Each datapoint represents an individual mouse; bars indicate mean values of each population. (**C**) Ratio of Tconv to Treg cells within 2W:I-A^b^ specific populations following 2W/CFA immunization of *CC10-USA* and *WT* littermate mice with or without prior *Smarta* transfer. Mean ± SEM values are shown (*n*=7-16 mice per group). Statistical significance was calculated via (**B**) unpaired t test or (**C**) one-way ANOVA with Tukey’s multiple comparisons test; ns, not significant, *p < 0.05, **p < 0.01.

Given these findings, we asked whether 2W:I-A^b^-specific Tregs were responsible for preventing 2W:I-A^b^-specific Tconv from proliferating following adoptive transfer of *Smarta* T cells into *CC10-USA* mice. To test the role of Tregs in our experimental system, we utilized the *Foxp3^DTR^* mouse model which enables inducible specific ablation of Tregs through the systemic administration of diphtheria toxin (DT) ^37^. CD4^+^ T cells from *Smarta* x *Foxp3^DTR^* mice (*Smarta-Foxp3^DTR^*) were adoptively transferred into *CC10-USA* x *Foxp3^DTR^* mice (*CC10-USA-Foxp3^DTR^*) and DT was administered every other day until harvest 7 days later (Fig. 4A). Surprisingly, despite effective ablation of all host and *Smarta* Tregs, there was no expansion, but rather a modest decrease of the 2W:I-A^b^-specific Tconv cells in the SLO and lungs (Fig. 4B). A similar lack of 2W:I-A^b^-specific Tconv expansion was observed when CD8^+^ T cells from *OT-I* mice were transferred into *CC10-USA-Foxp3^DTR^* mice (Fig. S3A). However, we did note an increase in overall CD4^+^ and CD8^+^ T cell infiltration within the lung parenchyma following ablation of endogenous host Tregs (Fig. S3B-C). These results demonstrate that 2W:I-A^b^-specific Tregs are not necessary to prevent activation of their Tconv counterparts in response to lung injury, but may play a role in reducing pathology by limiting trafficking of autoreactive T cells into the lung.

**Figure 4.**
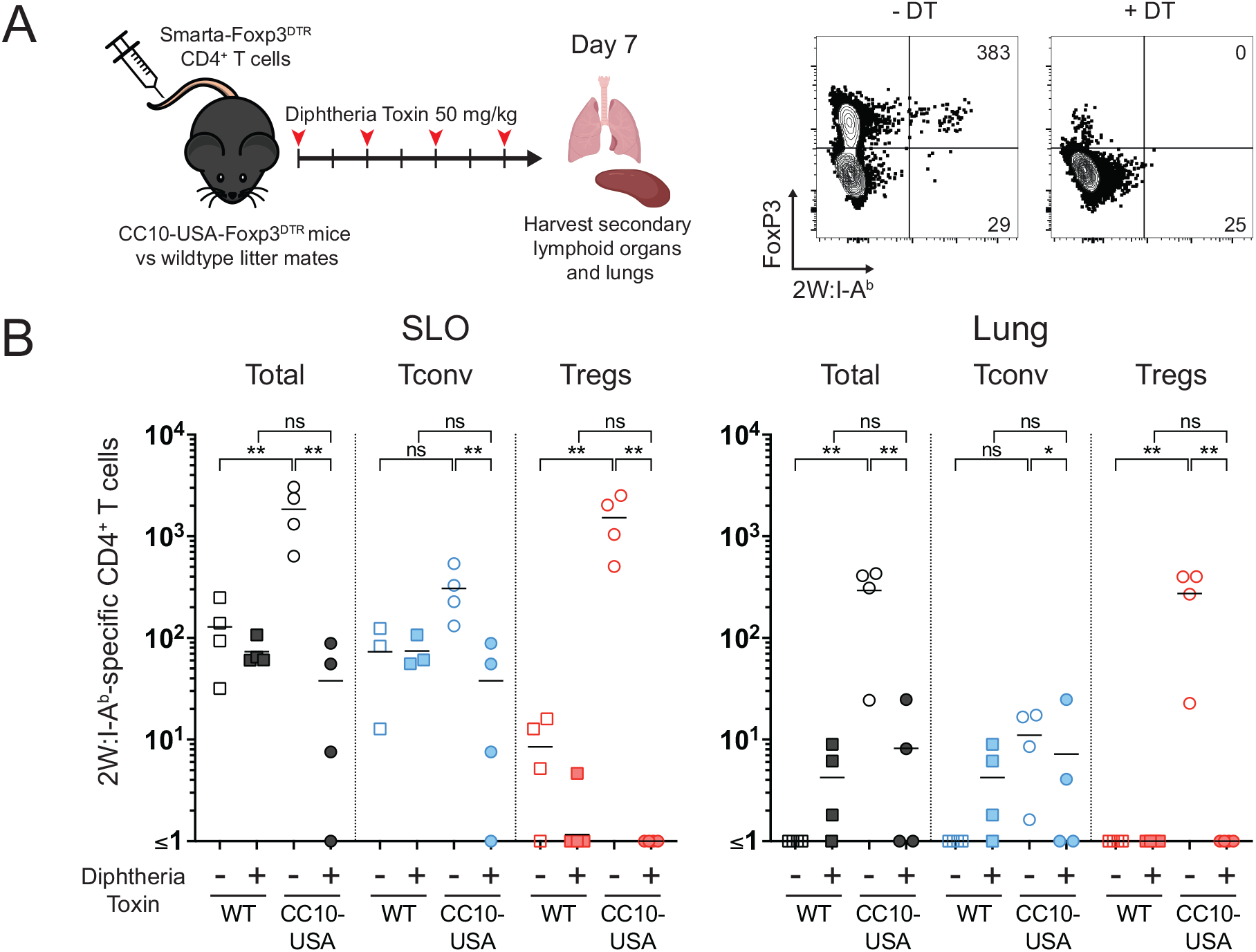
2W:I-A^b^-specific Tregs are not required for the regulation of 2W:I-A^b^-specific Tconv during immune-mediated tissue injury. (**A**) Experimental plan to test Treg function by global ablation. *CC10-USA-Foxp3^DTR^* or wildtype *Foxp3^DTR^* mice were treated with 50 mg/kg of diphtheria toxin (DT) *i.p*. every 2 days starting on the day of *Smarta-Foxp3^DTR^* CD4^+^ T cell transfer. Representative flow cytometry plots of CD4^+^ T cells in the lungs of *CC10-USA-Foxp3^DTR^* mice 7 days post-*Smarta-Foxp3^DTR^* transfer with or without DT treatment. Numbers inside the quadrants indicate total numbers of 2W:I-A^b^-specific CD4^+^ T cells calculated for the lungs of the whole mouse. (**B)** Quantification of 2W:I-A^b^-specific CD4^+^ T cells, further subdivided into Tconv and Treg cells, in the SLO and lungs of *CC10-USA-Foxp3^DTR^* or wildtype *Foxp3^DTR^* littermate mice 7 days after *Smarta-Foxp3^DTR^* transfer with or without DT treatment. Each datapoint represents an individual mouse; bars indicate mean values of each population. Statistical significance was calculated via one-way ANOVA with Tukey’s multiple comparison test; ns, not significant, *p < 0.05, **p < 0.01.

### Self antigen-specific Tregs asymmetrically expand in a mouse model of acute lung injury

Several studies have demonstrated the role of Tregs in mediating recovery from acute tissue injury, including recent studies highlighting effects that extend beyond immune cell suppression^38–40^. However, little is known about the specificity of these Tregs. To investigate whether we observed a similar expansion of 2W:I-A^b^-specific Tregs following acute lung injury, we treated the *CC10-USA* mice with intranasal administrations of *E.coli-derived* lipopolysaccharide (LPS) – a well-characterized mouse model of acute lung injury (ALI) marked by transient weight loss, increased inflammatory markers and protein in bronchoalveolar lavage (BAL) fluid, and leukocyte infiltration into the lungs^41,42^. Several studies using LPS-mediated ALI have demonstrated the importance of Tregs in mediating recovery from the injury^42,43^. To assess whether the pattern of self antigen-specific Treg expansion seen in our autoreactive T cell adoptive transfer model also occurs in a generalized tissue injury model without a targeted self antigen, we administered LPS intranasally into *CC10-USA* mice (Fig. 5A). As expected, the mice experienced a transient decrease in weight over the first 3-4 days followed by recovery over another 5-6 days (Fig. S4A-B). At 12 days post-treatment, we found a small but significant increase of 2W:I-A^b^-specific T cells in the lungs of *CC10-USA* but not *WT* mice (Fig. 5B). Compared to total lung infiltrating CD4^+^ T cells, the overwhelming majority of 2W:I-A^b^-specific T cells were again Foxp3^+^ Tregs (Fig. 5C).

**Figure 5.**
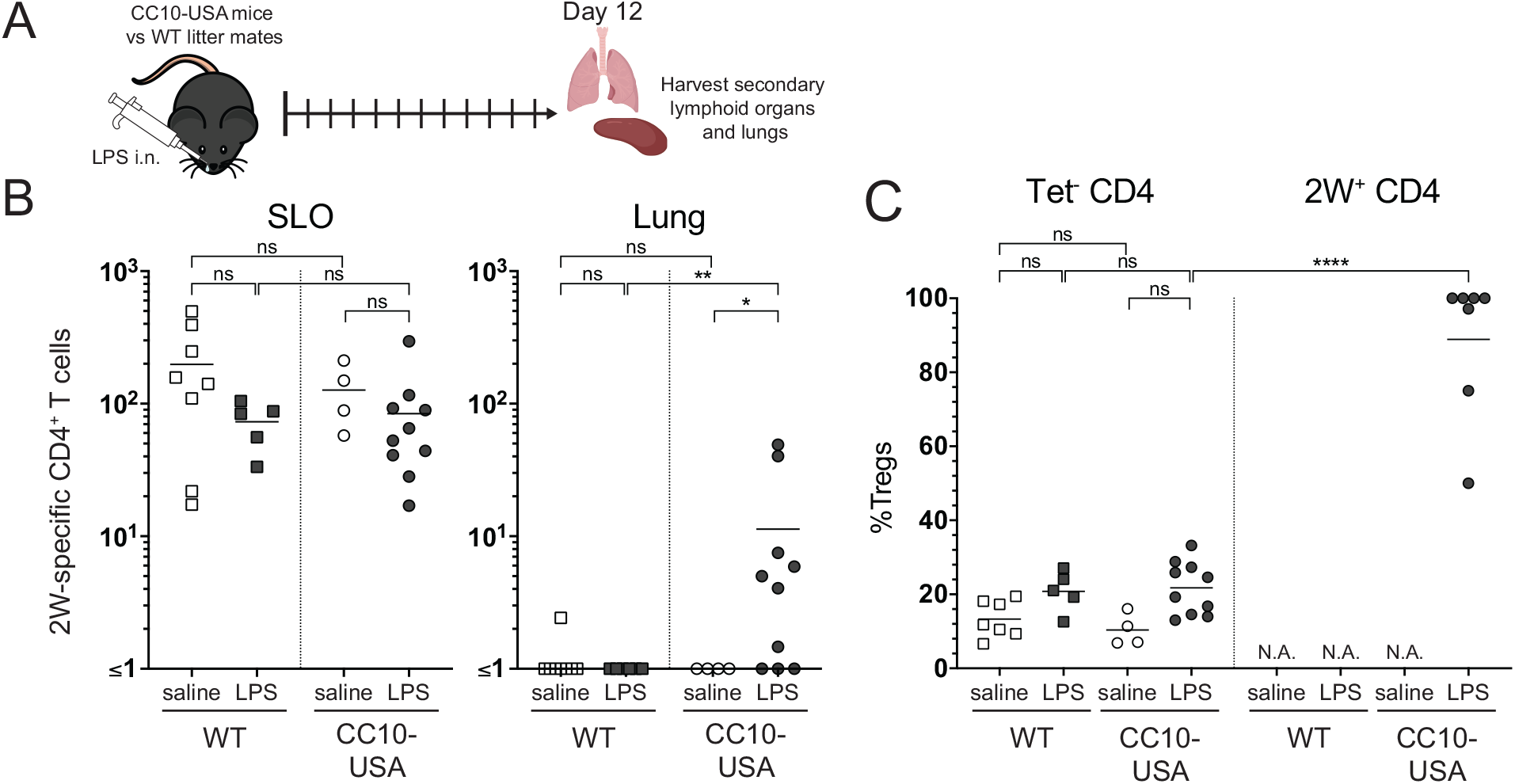
LPS-driven model of acute lung injury results in expansion of 2W-specific Tregs. (**A**) Experimental plan for intranasal (*i.n.*) LPS-mediated acute lung injury model. (**B**) Quantification of 2W:I-A^b^-specific CD4^+^ T cell numbers in the SLO and lungs 12 days following LPS treatment. Each datapoint represents an individual mouse; bars indicate mean values of each population. Statistical significance was calculated via Kruskal-Wallis test; ns, not significant, *p < 0.05, **p < 0.01. (**C**) Proportions of Foxp3^+^ Tregs within lung bulk CD4^+^ and 2W:I-A^b^-specific CD4^+^ T cell populations in *CC10-USA* and *WT* littermate mice 12 days following LPS treatment. Each datapoint represents an individual mouse; bars indicate mean values of each population. Statistical significance was calculated via one-way ANOVA with Tukey’s multiple comparison test; ns, not significant, ****p < 0.0001.

To further assess whether the Tregs that expand in the lungs following acute injury is a result of self antigen recognition, we leveraged the fact that all T cells from *Smarta* mice express a transgenic TCR that recognizes the gp66 antigen that is not present in mice and therefore the entire T cell repertoire lacks self antigen-specificity. We treated *Smarta* mice with LPS and compared them to *WT* mice as well as *RAG1-* deficient mice which are susceptible to LPS-mediated ALI due to their complete lack of Tregs^42^. *Smarta* mice experienced more severe weight loss and longer recovery times than *WT* controls at a level approaching that of *RAG1*-deficient mice (Fig. 6A). Furthermore, while LPS treatment of *Smarta* mice resulted in increased CD4^+^ T cell trafficking into the lungs compared to *WT* mice, there was a notable lack of Foxp3^+^ Tregs when compared to Tconv cells (Fig. 6B, S5A-B).

**Figure 6.**
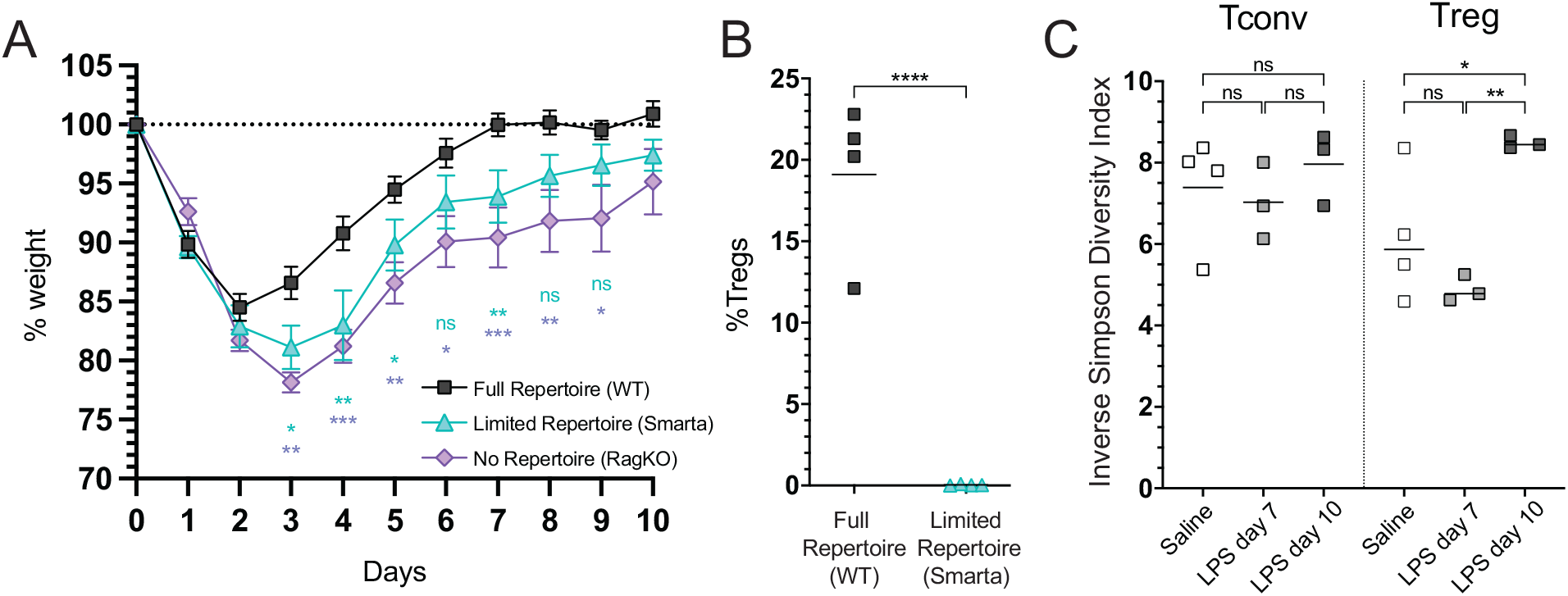
Self antigen-specificity is required for Treg development following LPS-mediated acute lung injury. (**A**) LPS was administered intranasally (*i.n.*) to *WT*, *Smarta*, or *Rag1KO* mice representing mice with a full, limited (no self antigen-specificity), and no T cell repertoire, respectively. Daily weights were monitored and plotted as relative values to baseline prior to treatment. Mean ± SEM are shown (*n*=4-7 mice per group). Statistical significance was calculated using a mixed-model (REML) with Tukey’s multiple comparison test; ns, not significant, *p < 0.05, **p < 0.01, ***p < 0.001. (**B**) Proportions of Foxp3^+^ Tregs within CD4^+^ T cells in the lungs of *WT* vs. *Smarta* mice 12 days following LPS treatment. Each datapoint represents an individual mouse; bars indicate mean values of each population. Statistical significance was calculated via unpaired t test; ****p < 0.0001. (**C**) TCR Vβ gene segment usage within lung-infiltrating CD4^+^ T cell populations of lungs was assessed by flow cytometry. Overall Vβ gene heterogeneity among Tconv and Treg populations at days 7 and 10 following LPS treatment was scored using the Inverse Simpson Diversity Index. Each data point represents an individual mouse; bars indicate mean values of each population. Statistical significance was calculated via one-way ANOVA with Tukey’s multiple comparison test; ns, not significant, *p < 0.05, **p < 0.01.

This was in stark contrast to the diversification of the lung Treg population in *WT* mice treated with LPS, as indicated by their increased usage of TCR variable gene segments over time that would reflect an increasing complexity of self antigen-specificities in the responding Treg repertoire (Fig. 6C). Collectively, our results suggest that Tregs expand upon recognition of self antigen during acute lung injury and play an important role in limiting further pathology.

## Discussion

The presence of abundant self antigen-specific T cells in the mature peripheral immune repertoire poses a constant threat for autoimmunity^5,44^. In their steady state, these T cells are heavily regulated by various mechanisms of tolerance that have been shown to limit priming and development into pathogenic autoreactive effector cells^2,4,28,45^. However, it is unclear how self antigen-specific T cell populations actually respond to recognition of self antigen that occurs in the context of acute inflammation and tissue damage that could precipitate an autoreactive immune response if normal tolerance mechanisms fail. Indeed, such long-term outcomes have been reported in viral-induced autoimmunity, cancer immunotherapy, and organ transplant rejection^19,46,47^. Moreover, EAE models demonstrate how such conditions can lead to epitope spreading of autoreactive T cell responses from one myelin sheath antigen to another, causing the relapsing-remitting nature of disease^22,24^.

In our current study, we directly tracked endogenous lung self antigen-specific CD4^+^ T cell populations to demonstrate that there is an initial T cell response to self antigen presentation during acute lung injury that is asymmetrically skewed towards the Treg compartment. The preferential activation of self antigen-specific Tregs was seen across different modalities of tissue damage, including both CD4^+^ and CD8^+^ T cell-mediated autoimmunity to neighboring self antigens, and an antigen-free model of LPS-mediated acute lung injury, suggesting that this is a generalizable response to self antigen presentation during inflammation.

The development of self antigen-specific Tregs during acute tissue damage has been implicated previously by several studies^38–40,48–50^. However, the self antigen specificity of Tregs in these studies has been circumstantial or inferred from the behavior of adoptively transferred naive TCR transgenic T cells serving as a surrogate population. By using a tetramer-based approach to directly study endogenous self antigen-specific T cells, our studies specifically demonstrated that self antigen-specific Tregs indeed expand and localize to the damaged tissue expressing the self antigen. This process was dependent on specific recognition of self antigen as the blocking of cognate peptide:MHC complex presentation to T cell receptors abrogated Treg development. Moreover, the expansion of self antigen-specific Tregs was dependent on signals from IL-2, but not TGFβ or IL-10. This is consistent with prior studies demonstrating that Tregs proliferate within lymph nodes in response to IL-2 produced by neighboring conventional T cells^51,52^. Within this framework, we propose that exclusive expansion of 2W:I-A^b^-specific Tregs in *CC10-USA* mice occurred through a process akin to “epitope spreading”, wherein the initial autoreactive *Smarta* or *OT-I* T cell responses against gp66 or OVA epitopes in the USA molecule led to the expansion of 2W:I-A^b^-specific Tregs. This may have also occurred in the LPS injury model, albeit the identities of other involved self antigens are unknown. In either case, these self antigenspecific Tregs expanded in the priming environment of secondary lymphoid organs in response to the IL-2 produced by co-localized autoreactive T cells. They then migrated to the site of tissue damage, the lungs, as evidenced by the lack of any appreciable amounts of pre-existing self antigen-specific Tregs in that tissue before injury. This process diverges from the usual connotation of “epitope spreading” in that the expanded Tregs do not promote further inflammation and disease, but rather limit it. Also noteworthy, the paracrine signaling of IL-2 from self antigen-specific effector T cells to self antigen-specific Tregs primed by the same antigen presenting cells offers a targeted mechanism that would increase the efficiency by which antigen-specific tolerance is achieved.

Accordingly, we did not detect Foxp3^+^ Tregs within the lungs of mice harboring a peripheral T cell repertoire with limited self antigen-specificity following intranasal LPS administration, and these mice experienced more weight loss than mice with normal repertoires. Additionally, mice treated with IL-2 blocking antibody or experiencing Treg ablation similarly fared worse after intranasal LPS administration (data not shown). These findings thereby suggest that the expansion and localization of Tregs at the site of tissue damage requires a T cell repertoire that harbors self-reactivity, and that these Tregs are important in regulating the ongoing inflammation. We can additionally infer that the infiltrating Tregs themselves are self antigen-specific in nature. This is consistent with the results of other studies in which transgenic T cells expressing the TCR of an expanded Treg from damaged skeletal muscle also localized to damaged skeletal muscle in other mice after adoptive transfer^39,53^. Collectively, these findings propose a model of immune regulation where the self antigen-specific T cell repertoire serves as a repository of Tregs that can readily expand in response to release of self antigens during acute tissue injury to limit further damage by pathogenic immune cells.

Expanded self antigen-specific Tregs were able to suppress 2W:I-A^b^-specific Tconv cells during subsequent immunization with peptide, but interestingly, the global ablation of Tregs did not lead to spontaneous activation and expansion of 2W:I-A^b^-specific Tconv populations during acute tissue injury. While this may seem to contradict our current model of steady state Treg-mediated tolerance of self antigen-specific T cells^2,28^, it is quite possible that the 2W:I-A^b^-specific Tconv cells are also under additional intrinsic mechanisms of regulation such as quiescence, anergy, or exhaustion, perhaps imprinted by their earlier continuous suppression from Tregs, a topic that remains to be explored. However, global Treg ablation has previously been shown to result in the rapid onset of systemic autoimmunity^37^, so other self antigenspecific Tconv cells are presumably not under the same level of restraint. Expanded 2W:I-A^b^-specific Tregs might contribute more broadly to the suppression of such cells.

Lastly, the mechanisms by which expanded self antigen-specific Tregs mediate their regulatory function remains to be elucidated. Moreover, recent studies investigating the expansion of Tregs in tissues during injury have suggested a novel, non-suppressive role in tissue repair, although the TCR specificities of these Tregs have only been indirectly linked to self antigens^38,39,53–55^. Further studies characterizing the phenotypic diversity of Tregs within the context of a self antigen-specific population will be important to elucidate the potential speciation of these Treg subsets.

By providing the most clear and direct demonstration to date of an endogenous self antigen-specific T cell population responding to acute tissue injury, we have revealed a strong propensity of self antigen-specific CD4^+^ T cells to polarize towards a regulatory fate. Our studies further suggest that this process is representative of a larger repertoire of self-reactive Tregs that expand in response to tissue injury to help regulate ongoing damage. Further studies into the mechanisms underlying the preferential expansion of these Tregs and the in-depth characterization of their effector function and targets promises to provide a broader understanding of the role of the self antigenspecific T cell repertoire in peripheral regulation, fresh insights underlying the loss of tolerance during the onset of autoimmunity, and potential novel therapeutic targets for modulating self-reactive Tregs in human disease.

## Materials and Methods

### Mice

Generation of the *USA* mouse has been previously described^28^. *Foxp3^RFP^*^56^, *Foxp3^DTR^*^37^, *Smarta*^57^, *OT-I*^58^, *Rag1KO*^59^, and *CD45.1*^60^ mice were purchased from Jackson Laboratory. *CC10-Cre*^61^ and *10BiT*^62^ mice were provided by T. Mariani (University of Rochester) and C. Weaver (University of Alabama at Birmingham) respectively. *CC10-Cre* mice were crossed with *USA, Foxp3^RFP^*, and *10BiT* mice to generate *CC10-USA* mice. *CC10-Cre* mice were crossed with *USA, Foxp3^DTR^*, and *10BiT* mice to generate *CC10-USA-Foxp3^DTR^* mice. *Smarta* mice were crossed with *Rag1KO* and *10BiT* mice, and then further bred with *Foxp3^RFP^* or *Foxp3^DTR^* mice. All strains and combinations thereof were bred and maintained under specific pathogen-free conditions. All offspring were screened for expression of relevant transgenes including leaky expression of *USA* by genotyping at Transnetyx. All experiments were approved by the Institutional Animal Care and Use Committee of Massachusetts General Hospital, an American Association for the Accreditation of Laboratory Animal Care (AAALAC)-accredited animal management program.

### Blocking antibodies

Monoclonal antibodies against IL-2 (clone: S4B6-1), TGFβ (clone: 1D11.16.8), IL-10 (clone: JES5-2A5), and their corresponding isotype control antibodies were purchased from BioXcell. The monoclonal antibody against 2W:I-A^b^ (clone: W6, gift from M. Jenkins) was purified from the supernatants of cultured hybridoma cells. Where indicated, blocking antibodies were administered at 500 μg per mouse intraperitoneally (*i.p.*) on the day of adoptive transfer of T cells.

### Peptide:MHC tetramers

The generation of peptide:MHCII tetramers has been previously described in detail^63^. In brief, soluble heterodimeric I-A^b^ molecules covalently linked to 2W (EAWGALANWAVDSA) or LCMV gp_66-77_ (DIYKGVYQFKSV) peptide epitopes were expressed and biotinylated in stably transfected *Drosophila* S2 cells. Following immunoaffinity purification, these biotinylated peptide:MHCII complexes were titrated and tetramerized to R-phycoerythrin (PE), allophycocyanine (APC), or cyan-7 conjugated R-phycoerythrin (PECy7) fluorochrome-conjugated streptavidin (Prozyme, Thermo-Fisher).

### Adoptive T cell transfer

Spleens and lymph nodes were harvested from 6-12 week old male and female CD45.1-congenically marked donor *Smarta* or *OT-I* mice. CD4^+^ and CD8^+^ T cells were isolated using from *Smarta* or *OT-I* mice respectively, using magnetic bead-based negative selection according to manufacturer’s guidelines (Miltenyi). Purity of the transgenic TCRs were verified by flow cytometry staining for TCRVα2. The equivalent of 3×10^6^ *Smarta* or *OT-I* cells were injected retro-orbitally into 6-12 week old *CC10-USA* or *WT* littermates. Lungs, spleens, and lymph nodes were harvested and analyzed 7 days post-transfer. Where indicated, Foxp3^+^ Tregs were ablated in host mice via *i.p*. administration of diphtheria toxin (List Biologicals) at 50 mg/kg on the day of adoptive transfer and every other day afterwards until harvest.

### Immunizations

2W (EAWGALANWAVDSA) peptide was custom ordered from Genscript. 100 μg of peptide was prepared in a 50 μL emulsification of 1:1 (v/v) normal saline and complete Freund’s adjuvant (CFA)(Sigma) and injected subcutaneously at the base of the tail. For LPS treatments, 25-50 μg of E.coli derived lipopolysaccharide (List Biologicals) was prepared in 50 μL of normal saline and administered intranasally into anesthetized mice. Peptide/CFA immunized or LPS-treated mice were harvested for tissues 12 days posttreatment unless indicated otherwise.

### Tetramer-based cell enrichment and flow cytometry

Spleen and lymph nodes (SLO) were pooled and manually separated into single cell suspensions. Lung tissues were homogenized with a GentleMACS dissociator (Miltenyi) and digested with a cocktail of Liberase (70 μg/mL, Roche) and aminoguanindine (10mM, Sigma). Single cell suspensions of SLO or lungs were treated for 5 minutes with 50 nM Dasatinib (Sigma) at room temperature (RT) followed by staining with a premixed cocktail of the indicated tetramers at a final concentration of 10 nM for 1 hour at RT in the presence of anti-mouse CD16/32 antibody (clone 93, BioLegend). SLO preparations were magnetically enriched for tetramer-specific cells as previously described^64^ while lung samples were magnetically enriched for CD90.2 expression (Miltenyi). Lung resident cells were distinguished from those in the vasculature by exclusion from staining with fluorescently labeled anti-CD45 antibody (PerCP, clone 30F-11, BioLegend) that was injected retro-orbitally into each mouse approximately 3 minutes prior to euthanasia^65^. Magnetically enriched cells were stained with Fixable Viability Dye eFluor780 (eBioscience) and the following panel of fluorescently-labeled antibodies at 4C for 30 minutes: anti-CD90.2 (BUV395, clone 53-2.1, BD Biosciences), anti-CD90.1 (BUV737, clone OX-7, BD Biosciences), anti-CD45.1 (PacBlue, clone A20, BioLegend), anti-CD4 (BV421, clone GK1.5, BioLegend), anti-CD8 (PacOrange, clone 5H10, Invitrogen), anti-PD-1 (PerCP/Cy5.5, clone 29F.1A12, BioLegend), anti-CD11b (PE/Cy5, clone M1/70, BioLegend), anti-CD11c (PE/Cy5, clone N418, BioLegend), anti-F4/80 (PE/Cy5, clone BM8, BioLegend), anti-CD45R/B220 (PE/Cy5, clone RA3-6B2, BioLegend), and anti-CD44 (AlexaFluor700, clone IM7, BioLegend). Flow cytometry was performed on the Aurora (Cytek), LSRFortessa X-20 (BDIS), or Cytoflex S (Beckman Coulter) and analyzed by FlowJo software (TreeStar).

### Mouse IgM Enzyme-Linked Immunosorbent Assay (ELISA)

*CC10-USA* mice or *WT* littermates were euthanized, cannulated, and the bronchoalveolar lavage (BAL) fluid was collected in two 0.5-mL lavages with cold 600 uM EDTA in PBS. Intra-alveolar cells were removed from the BAL fluid via centrifugation. 96-well microtiter plates (Greiner) were coated overnight at RT with goat anti-mouse IgM affinity-purified polyclonal antibody (0.6 μg/well, Bethyl). Plates were blocked with 3% (w/v) bovine sera albumin in PBS for 45 minutes at RT before BAL samples were added to their respective wells and incubated for another 2 hours at RT. The wells were then washed with PBS/0.1% (v/v) Tween20, and HRP-conjugated goat anti-mouse IgM detection antibody (0.02 μg/mL, Bethyl) was added to the wells for 1 hour at RT prior to developing with 3,3’,5,5’-tetramethylbenzidine (TMB) substrate solution (BioLegend). The reaction was stopped by adding 0.1M sulfuric acid and the optical density was determined at 450 nm using a SpectraMax iD5 spectrophotometer.

### Confocal Immunofluorescence Microscopy

Lungs of *CC10-USA* mice or *WT* littermates were perfused with 2 mL of PBS + 2% (v/v) paraformaldehyde (PFA), harvested, and fixed in 2% PFA solution for 1 hour at 4C. The lung tissues were then placed sequentially into 10%, 20%, and 30% (w/v) sucrose PBS solutions prior to embedding into Optimum Cutting Temperature (O.C.T.) compound (Tissue-Tek) and frozen over dry ice. A Cryostat (Microm HM 505 E, GMI) was used to generate 20 μm sections which were then stored at −20C. Thawed slides were rehydrated in PBS at RT, permealized with 1% (v/v) Triton X-100 (Sigma), and blocked with Image-iT FX Signal Enhancer (Cell Signal) followed by PBS containing 0.1% (v/v) Tween20 and 5% (v/v) normal goat serum (Jackson Immunoresearch). Samples were then stained overnight at 4C with anti-CD4 (AlexaFluor488, clone RM4-5, BioLegend) and anti-CD45.1 (Alexafluor647, clone A20, BioLegend). Slides were then mounted with ProLong Diamond Antifade Mountant (Thermo Fisher). Immunofluorescence images were obtained on a Zeiss LSM780 confocal laser scanning microscope with a 1-Airy unit pinhole. Sequential scans with 1 laser line active per scan were used with excitation wavelengths of 405, 488, 561, and 633 nm; the multispectral GaAsP detector emission ranges were set to 410-513, 491-616, 585-733, 638-755 nm, respectively. Images were analyzed and overlays were generated with ImageJ v.2.1.0.

### Statistical Analysis

No statistical methods were used to predetermine sample sizes. All statistical analysis was performed on Prism (GraphPad). Specific tests utilized are indicated in figure legends. For visualization purposes, values equal to 0 were adjusted to 1 to appear on the axis of log-scaled data.

## Author Contributions

Conceptualization: D.S.S. and J.J.M.; Methodology: D.S.S. and J.J.M.; Investigation: D.S.S., S.R., C.N.C., and L.F.K.; Resources: R.A.R. and R.M.A.; Formal Analysis: D.S.S. and J.J.M.; Writing – Original Draft, D.S.S. and J.J.M.; Funding Acquisition: D.S.S. and J.J.M.; Supervision: J.J.M.

## Acknowledgments

We thank O.L. Venezia and J.M. Neary for assistance with mice and tetramers, J.W. Griffith and H. Amatullah for advice on BAL studies, R.W. Nelson for advice on mice immunizations, S. Banerjee and C.P. Phelan for technical assistance, and M.C. Takenaka and K.L. Jeffrey for constructive input. This work was funded by the National Institutes of Health (R01AI107020 and R01DK126910 to J.J.M., T32AI007512 to D.S.S.) and MGH ECOR (J.J.M.).

