## supplemental figures for "Lung injury induces a polarized immune response by self antigen-specific Foxp3^+^ regulatory T cells"

### Figure S1

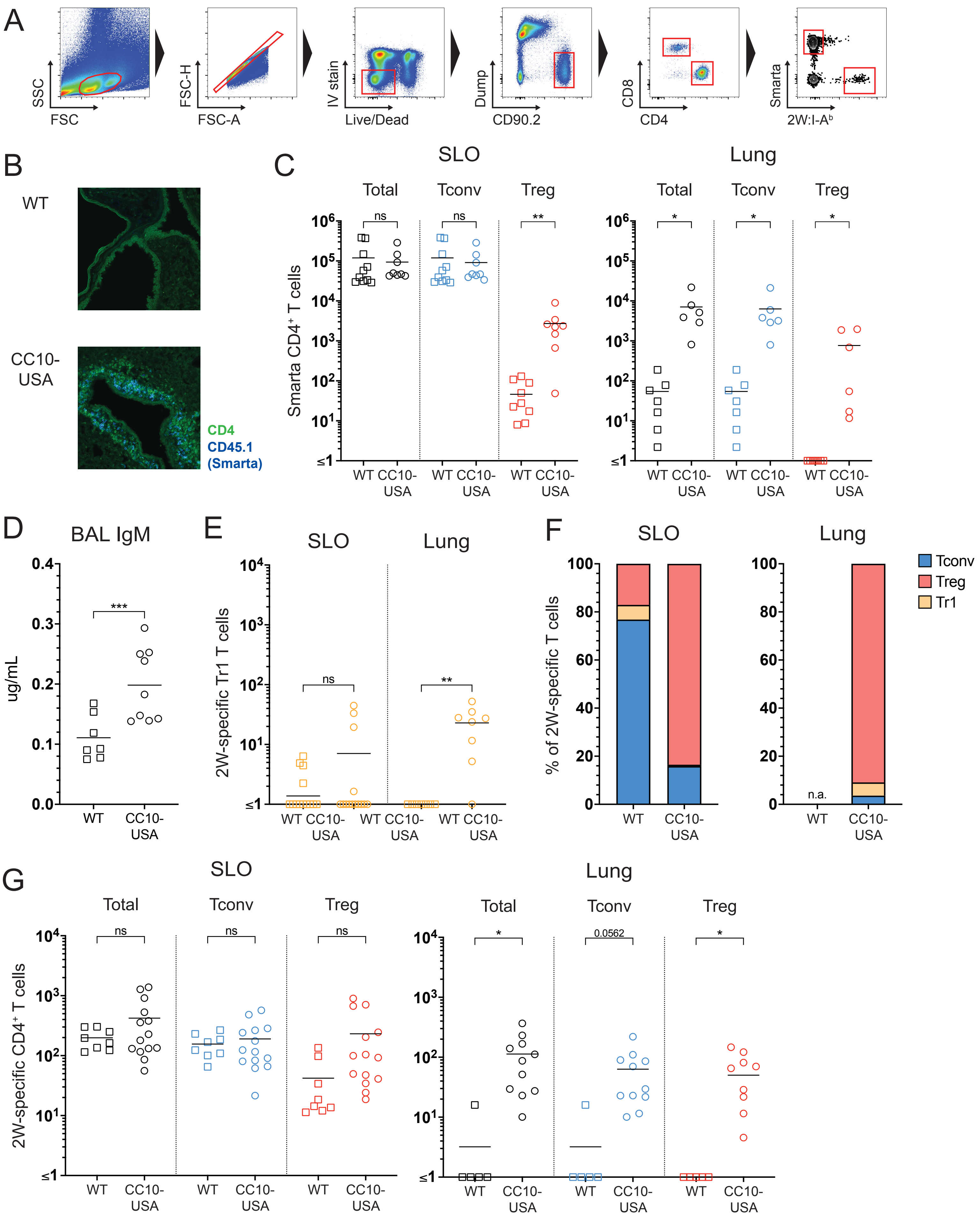
A
SSC
FSC
FSC-H
FSC-A
IV stain
Live/Dead
Dump
CD90.2
CD8
CD4
Smarta
2W:I-A^b
B
WT
CC10-USA
CD4
CD45.1
(Smarta)
C
SLO
Lung
Total
Tconv
Treg
Smarta CD4+ T cells
ns
**
*
WT
CC10-USA
D
BAL IgM
ug/mL
***
E
2W-specific Tr1 T cells
F
% of 2W-specific T cells
Tconv
Treg
Tr1
n.a.
G
2W-specific CD4+ T cells
0.0562

**Supplemental Figure 1. Transfer of USA-specific T cells into *CC10-USA* mice leads to tissue injury and asymmetric expansion of 2W:I-A$^b$-specific Tregs.**

(**A**) Gating strategy for identification of antigen-specific T cells via flow cytometry. Representative flow cytometry illustrating sequential inclusion and exclusion gates used to analyze 2W:I-A$^b$-specific or *Smarta* CD4$^+$ T cells following magnetic bead-based enrichment of SLO or lungs.

(**B**) Detection of *Smarta* CD4$^+$ T cells along the lung airways of *CC10-USA* mice on confocal immunofluorescence 7 days after transfer.

(**C**) Quantitative summary of *Smarta* CD4$^+$ T cells recovered from the SLO (left) and lungs (right) of *CC10-USA* mice and *WT* littermates 7 days post-transfer.

(**D**) Quantitation of IgM present in the BAL of *CC10-USA* and *WT* littermates 7 days after transfer of *Smarta* CD4+ T cells.

(**E**) Quantification of total 2W:I-A$^b$-specific Tr1 cells in the SLO (left) and lungs (right) of *CC10-USA* mice and *WT* littermates 7 days post-transfer.

(**F**) Relative proportions of 2W:I-A$^b$-specific T cells phenotypes in the SLO (left) and lungs (right) of *CC10-USA* mice and *WT* littermates 7 days post-transfer.

(**G**) Quantification of total 2W:I-A$^b$-specific CD4$^+$ T cells, subdivided into Tconvs and Tregs, of SLO and lungs from *CC10-USA* mice and *WT* littermates 7 days after adoptive transfer of OVA-specific CD8$^+$ T cells from *OT-I* mice.

Each data point represents an individual mouse. Bars indicate mean values of each population. Statistical significance was calculated via unpaired t test; n.a., not applicable, ns, not significant, *$p < 0.05$, **$p < 0.01$, ***$p < 0.001$, ****$p < 0.0001$.

**Figure S2**

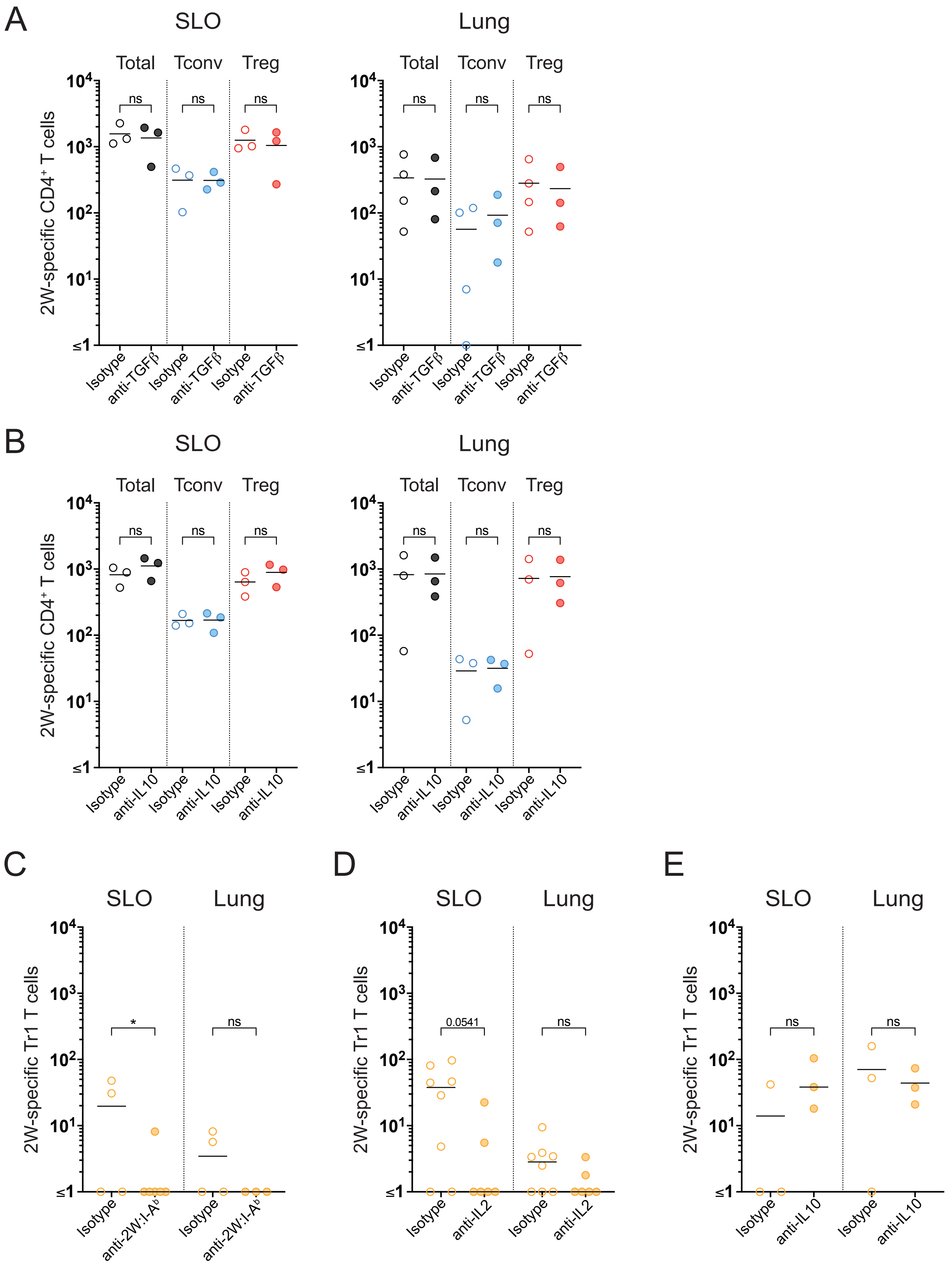

A
SLO
Lung
Total
Tconv
Treg
ns
2W-specific CD4+ T cells
Isotype
anti-TGFβ
B
anti-IL10
C
D
E
2W-specific Tr1 T cells
*
0.0541
anti-2W:I-Ab
anti-IL2

**Supplemental Figure 2. Blockade of TGFβ and IL-10 signaling does not abrogate 2W:I-A$^b$-specific Foxp3$^+$ Tregs expansion following transfer of *Smarta* CD4$^+$ T cells.**
(**A-B**) Quantification of total 2W:I-A$^b$-specific CD4$^+$ T cells, further subdivided into Tconvs and Tregs, in SLO (left) and lungs (right) of *CC10-USA* mice and *WT* littermates 7 days following *i.p.* administration of 500 μg of (**A**) anti-TFGβ or (**B**) anti-IL10 blocking antibodies versus isotype control at the time of adoptive transfer of *Smarta* CD4$^+$ T cells.
(**C-E**) Quantification of 2W:I-A$^b$-specific Tr1 cells in the SLO (left) and lungs (right) of *CC10-USA* mice and *WT* littermates 7 days after *i.p.* administration of (**C**) anti-2W:I-A$^b$, (**D**) anti-IL2, or (**E**) anti-IL10 blocking antibodies versus isotype controls at time of *Smarta* CD4$^+$ T cells transfer.
Each data point represents an individual mouse. Bars indicate mean values of each population. Statistical significance was calculated via unpaired t test; ns, not significant, $*p < 0.05$.

### Figure S3

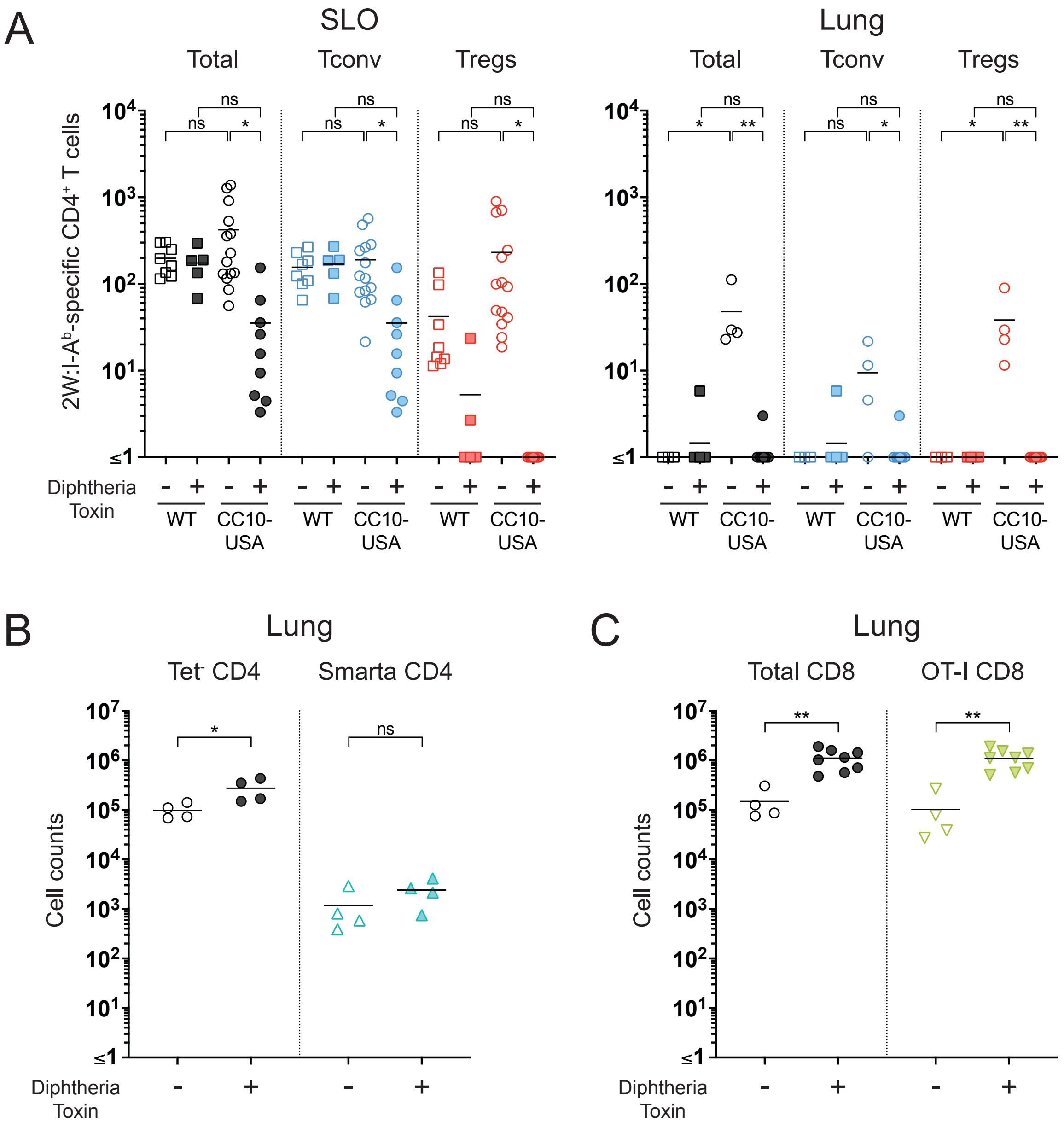


**Supplemental Figure 3. Depletion of endogenous Tregs leads to increased trafficking of CD4⁺ and CD8⁺ T cells into the lungs of *CC10-USA* mice following transfer of USA-specific T cells.**

(**A**) Quantification of total 2W:I-A$^b$-specific CD4$^+$ T cells, subdivided into Tconvs and Tregs, following tetramer-based cell enrichment of SLO (left) and lungs (right) from *CC10-USA-Foxp3$^{DTR}$* mice or wildtype *Foxp3$^{DTR}$* littermates 7 days after adoptive transfer of *OT-I* CD8$^+$ T cells with or without DT treatment.

(**B**) Quantification of tetramer negative (left) and *Smarta* (right) CD4$^+$ T cells infiltrating into the lungs of *CC10-USA* mice and *WT* littermates 7 days after adoptive transfer of Smarta CD4$^+$ T cells with or without DT treatment.

(***C***) Quantification of total (left) and *OT-I* (right) CD8$^+$ T cells infiltrating into the lungs of *CC10-USA mice* and *WT* littermates 7 days after adoptive transfer of *OT-I* CD8$^+$ T cells with or without DT treatment.

Each data point represents an individual mouse. Bars indicate mean values of each population. Statistical significance was calculated via (**A**) one-way ANOVA with Tukey's multiple comparisons test or (**B**-**C**) unpaired t test; ns, not significant, *$p < 0.05$, **$p < 0.01$.

**Figure S4**

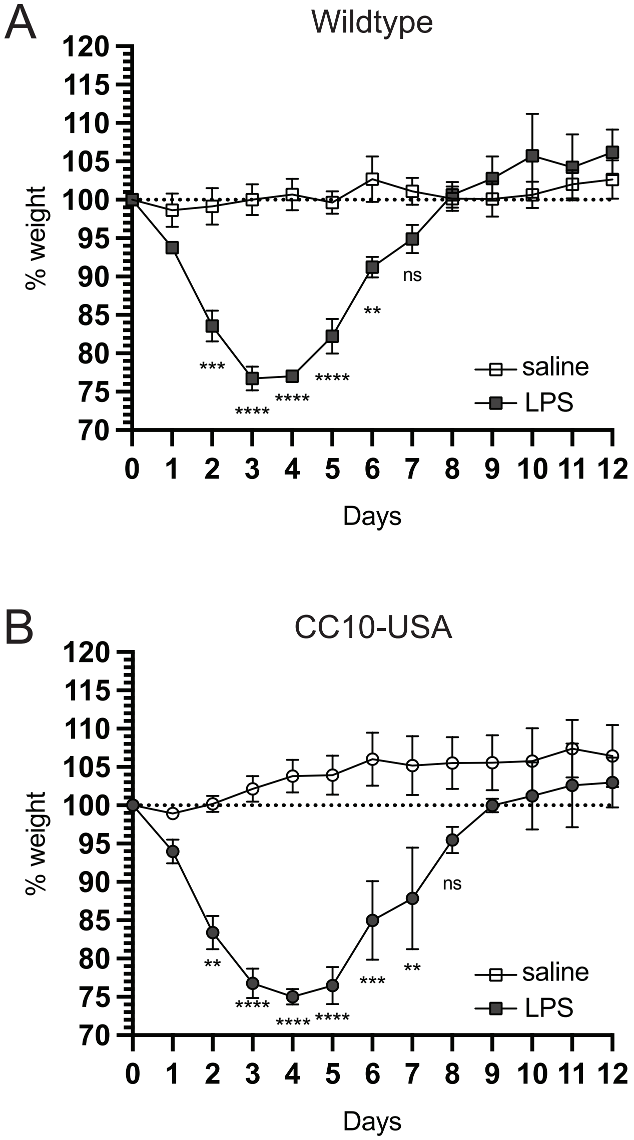


**Supplemental Figure 4. Weight loss following intranasal LPS treatment.**
(**A-B**) Intranasal LPS was administered to (**A**) *WT* or (**B**) *CC10-USA* mice and daily weights were monitored. Changes in weight were plotted relative to baseline prior to treatment. Mean ± SEM are shown (*n*=3-4 mice per group). Statistical significance was calculated using a RM 2-way ANOVA with Šidák's multiple comparison test; ns, not significant, **p < 0.01, ***p < 0.001, ****p < 0.0001.

**Figure S5**

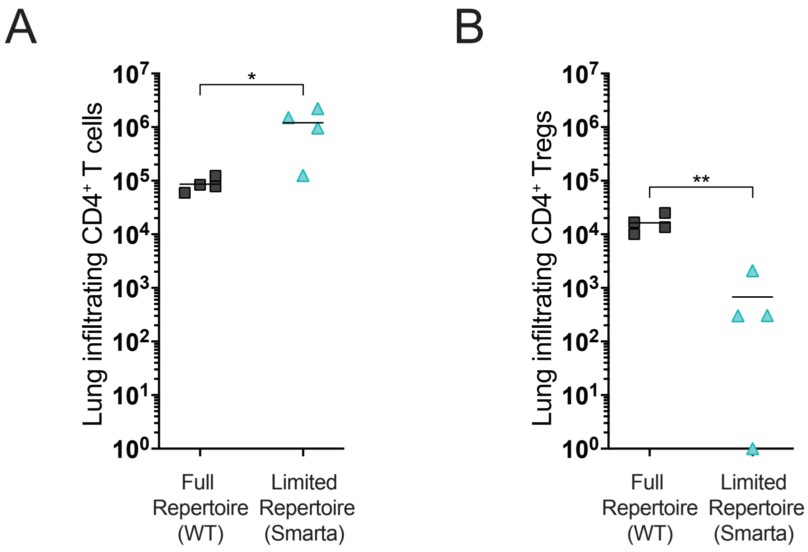


**Supplemental Figure 5. Decreased infiltration of Tregs in mice with limited self antigen-specific T cell repertoire.**
(A-B) Quantification of (A) total $CD4^+$ T or (B) Tregs cells infiltrating into the lungs of *Smarta* mice vs *WT* littermates 10 days after intranasal LPS treatment. Each data point represents an individual mouse. Bars indicate mean values of each population. Statistical significance was calculated via unpaired t test; $*p < 0.05$, $**p < 0.01$.
